# Hue tuning curves in V4 change with visual context

**DOI:** 10.1101/780478

**Authors:** Ari S. Benjamin, Pavan Ramkumar, Hugo Fernandes, Matthew Smith, Konrad P. Kording

**Author notes:** Correspondence, Richards 401A 3700 Hamilton Walk. Philadelphia PA, 19104.

## Abstract

Neurons are often probed by presenting a set of stimuli that vary along one dimension (e.g. color) and quantifying how this stimulus property affect neural activity. An open question, in particular where higher-level areas are involved, is how much tuning measured with one stimulus set reveals about tuning to a new set. Here we ask this question by estimating tuning to hue in macaque V4 from a set of natural scenes and a set of simple color stimuli. We found that hue tuning was strong in each dataset but was not correlated across the datasets, a finding expected if neurons have strong mixed selectivity. We also show how such mixed selectivity may be useful for transmitting information about multiple dimensions of the world. Our finding suggest that tuning in higher visual areas measured with simple stimuli may thus not generalize to naturalistic stimuli.

**New & Noteworthy:** Visual cortex is often investigated by mapping neural tuning to variables selected by the researcher such as color. How much does this approach tell us a neuron’s general ‘role’ in vision? Here we show that for strongly hue-tuned neurons in V4, estimating hue tuning from artificial stimuli does not reveal the hue tuning in the context of natural scenes. We show how models of optimal information processing suggest that such mixed selectivity maximizes information transmission.

## Introduction

Neurophysiology has long investigated the visual cortex by asking how its various areas encode the visual world. In order to simplify experimental design, the majority of early work constructed stimuli sets in which only a few key visual parameters varied. By then observing which areas show corresponding changes in neural activity, the visual cortex can be described in terms of the variables for which neurons are highly tuned. This program has been successful in characterizing that the response properties of neurons in the ventral stream ascend in complexity. V1 is discussed as responding to “edge-detecting” Gabor filters [1], V2 to variations in local curvature [2], V4 to more complex shapes [3], and IT to specific objects and faces [4], which together have inspired the theory that object recognition proceeds via hierarchical image representations [5–7].

In recent decades there has been a greater interest in understanding how much the response to simple, parameterized stimuli is informative about how the cortex encodes naturalistic stimuli of the type that make up everyday experience. In addition to representing a field-wide shift towards ethology and natural paradigms, this question pertains to how much tuning curves estimated from artificial stimuli are good models of neurons’ general functional role in visual processing. If tuning varies widely and unpredictably with context, the knowledge gained from simplified stimuli would be unique and particular to the tested stimuli alone. It is therefore critical to test how much tuning inferred from simplified stimuli is informative of tuning for more complicated stimuli.

In principle, such a question could be asked by steadily increasing the number of stimulus parameters that are varied until the complexity of stimuli approaches that of natural scenes. The original tuning curve could then be appreciated in the context of all others, and interactions with other variables characterized. However, this approach is prohibitive because the number of stimuli required to be displayed to an animal scales exponentially with the number of parameters separately varied. Any practical experiment of this type would need to leave many potential parameters unvaried.

An alternative paradigm, and one that has a long history in early visual areas, is to characterize neurons directly from their responses to natural images by regressing models of the visual encoding [8–13]. To build models without a number of stimuli exponential in their dimension, one must place constraints upon what visual encodings are viewed as possible. For early areas scientists often fit rather simple encoding models, such as linear or linear-nonlinear models. Studies in this vein have found that many aspects, of the V1 response, such as preferred orientation, appear similar between both natural images and drifting gratings [11]. However, other aspects of the V1 response are different for natural images [10], which limits how well artificial stimuli responses can inform researchers about natural stimuli responses [12, 14]. In area V1, many tuning properties are shared between responses to artificial and natural stimuli, raising the question if the same is true for higher level areas.

For higher areas, however, a regression methodology has not yet allowed such a comparison. In this work we focus on area V4. While the response of V4 to natural scenes been studied [5, 15–18], most knowledge about the tuning of V4 has derived from parameterized stimuli sets and it is not known how much tuning can vary with context. Our specific focus is hue, as V4 was first characterized as a color area [19] before later studies found selectivity for other visual features (such as orientation [20], curvature [3], shape [21–23], depth [24–26], and motion [27]; reviewed in [28]). As revealed by displaying simple colored shapes, color-selective neurons are predominantly located in color ‘globs’ [29], which intersperse ‘interglob’ regions more selective for orientation [30]. Glob cells are further arranged by their color preference [31] and generally in the same hue sequence as is found in perceptual color space [32, 33]. It is not known how accurately these tuning curves describe V4’s response to color in natural stimuli.

In this work, we tested the limits of how much tuning in macaque V4 generalizes to complex stimuli by estimating the hue tuning curves of color-responsive neurons by varying the hue of simple stimuli, and then again from their responses to natural scenes. That is, we asked how well P(Y|X, Z=z), which is the probability of spike counts Y given hue X and a fixed context z, stands in for P(Y|X), the average hue tuning over natural images. We regressed hue tuning from natural scene responses using a progression of encoding models, including one based on a deep artificial network pretrained to classify images [5], obtaining a similar conclusion from each. Overall we found that, although hue strongly modulates the V4 response, the tuning curves estimated from responses to stimuli of a single hue poorly described how hue affected responses to natural scenes. Lastly, we show how such mixed selectivity may be useful for information transmission.

## Materials and Methods

### Experimental setup: recordings

We recorded from 96-electrode Utah arrays (1.0 mm electrode length) implanted in visual area V4. At the time of the experiment, Monkey 1 (M1) was aged 5 years, 10 months and Monkey 2 (M2) was aged 9 years, 4 months. Surgical details describing the implantation method can be found in previous publications [34, 35]. The array was located in the left hemisphere for monkey M1 and in the right hemisphere for M2. Spikes were sorted off-line first with an automated clustering procedure [36] and then refined by hand using custom MATLAB software (https://github.com/smithlabvision/spikesort) taking into account waveform shape and interspike interval distributions [37].

All experimental procedures were approved by the Institutional Animal Care and Use Committee of the University of Pittsburgh.

### Artificial stimuli

Both monkeys viewed uniform images of a single hue on a computer screen at 36 cm distance, with a resolution of 1024×768 pixels and a refresh rate of 100 Hz on a 21” cathode ray tube display. We found that full-field hues elicited strong and selective responses from a majority of neurons (see Results). The hues were sampled from the hue wheel in CIELUV color space (calculated with a D65 standard illuminant and standard observer) at increments of 1 degree and at a chromaticity ensured to lie in the RGB gamut, and were presented in random sequence. Monkey M1 freely viewed the stimuli, and was rewarded periodically for maintaining eye position on the screen for 4 seconds, after which time the static image was refreshed. The trial was ended if the monkey looked beyond the screen during this duration. Monkey M2 was trained to fixate a small dot at the center of the screen for 0.3 seconds, during which three images were flashed for 100ms each. A 0.5 second blank period interspersed each fixation. Monkey 1 viewed 7,173 samples of the uniform hue stimuli over 10 sessions, while Monkey 2 viewed 1,119 samples during a single session. The full monitor subtended 55.5 degrees of visual angle horizontally and 43.1 degrees vertically. The monitor was calibrated to linearize the relationship between input luminance and output voltage using a lookup table. This calibration was performed for grayscale images, and the color profile of the monitor was not separately calibrated.

### Natural images

Both monkeys viewed samples from a dataset of 551 natural images, obtained from a custom-made Google Images web crawler that searched and downloaded images based on keywords such as cities, animals, birds, buildings, sports, etc. Monkey M1 viewed images over 15 separate sessions, for a total of 77961 fixations. Monkey M2 viewed images over two sessions on a single day, for a total of 6713 fixations. We then extracted the features from the image patch centered around each fixation that would serve as model inputs. The image patch around fixation corresponded to the 400 x 400 pixel block surrounding the center of gaze. This corresponds to a region 23.5 visual degrees on a side.

### Gaze tracking and fixation segmentation

We employed a free-viewing paradigm for one monkey (M1) and a fixed-gaze paradigm for the other (M2). The location of each monkey’s gaze on the screen was tracked with an Eyelink 1000 infrared tracker (SR Research, Ottawa, Ontario, Canada). Visual stimuli were presented and the experimental trials were controlled by custom MATLAB software in conjunction with the Psychophysics Toolbox [38]. For monkey M1, we segmented each fixation as a separate event based on thresholding the position and velocity of the gaze coordinates. We did not analyze activity occurring during eye movements. Once each fixation was separated, the average location of the fixation was recorded and matched to image coordinates. Monkey M2 was trained to fixate on a dot positioned at the center of each image. The gaze was tracked as for M1, but this time only to enforce fixation and terminate the trial if the gaze shifted away from center.

### Session concatenation

Although all recordings in M1 were performed with the same implanted Utah array, they were recorded over several sessions. The recordings for M2 were made in a single session. In M1, this introduced the possibility that the array might have drifted, and that a single channel might have recorded separate neurons in different sessions. To address this possibility, we noted that spikes identified in a channel in one session will be less predictive of another session’s activity if the neurons are not the same, as we expect tuning to be relatively static across days [39, 40]. We thus filtered out neurons whose uniform hue tuning changed across sessions. We trained a gradient boosting regression model with Poisson targets to predict spike counts in response to the hue of the stimuli. Nuisance parameters, such as duration of stimulus, gaze position, inter-trial interval, etc., were also included as model covariates to increase the predictive power even for neurons that were not hue-tuned. We then labeled a neuron as having static tuning as follows. First, we trained the model on each single session in a 8-fold cross-validation procedure and recorded the mean pseudo-R^2^ score. This score reflected how well the model could predict held-out trials on the same session. Then, we re-trained the model on each session and predicted on a different session, for all pairs of sessions. This resulted in a cross-prediction matrix with diagonal terms representing same session predictability (the 8-fold CV score), and off-diagonal terms representing generalization between sessions. We did not concatenate sessions if hue tuning estimated in one session could not predict hue responses in another session (i.e. the CV pseudo-R^2^ score was less than 0).

The natural image sessions were interspersed with the artificial sessions. If a natural image session occurred between two artificial sessions, and a neuron showed static tuning both artificial sessions as identified in the above manner, then that natural image session was included for the hue tuning comparison and model fitting. The recordings of units from other natural image sessions were not used. This procedure improved our confidence that the neurons recorded in different sessions were the same.

### Uniform hue tuning curve estimation

Hue tuning curves were built for each neuron by plotting its spike rate on each fixation against the observed hue. Spike rates were calculated from activity 50ms after fixation onset until 300ms or fixation offset, whichever came first. For the visualizations in the figures, we performed LOWESS smoothing, in which each point of the curve is given by a locally-weighted linear regression model of a fraction of the data. The error envelope of the curve represents the 95% confidence interval given by bootstrapping over individual fixations. To calculate the correlation between tuning curves, we did not correlate the LOWESS-smoothed curves but rather the simple binned averages. We created 16 bins of hues and calculated the average spike rate for all stimulus presentations of those hues, then correlated the 16-dimensional tuning curve vector with the natural image tuning curves.

To see how well simple tuning could explain the V4 response, we interpreted these tuning curves as models of the natural image (presented in Results in Figure 2B). This was a linear model whose coefficients are set from the uniform field tuning curve. Prediction was performed such that an image patch that was all a single color would result in a prediction that was the firing rate observed in the uniform field condition, and mixtures of colors would predict linear combinations of the corresponding observed firing rates. More precisely, the predicted firing rate was a dot product of the tuning curve with the (normalized) hue histogram. To extract these hue histograms from image patches, we calculated the hue angle of each pixel in the receptive field during a given stimulus presentation (see Receptive Field estimation below) in CIELUV space. We then binned these hues into histograms with 16 bins of hues, and these histograms served as the representation of hue input to the model. The final predicted response was then added to a constant term to account for the difference in mean firing rate across contexts.

**Figure 2.**
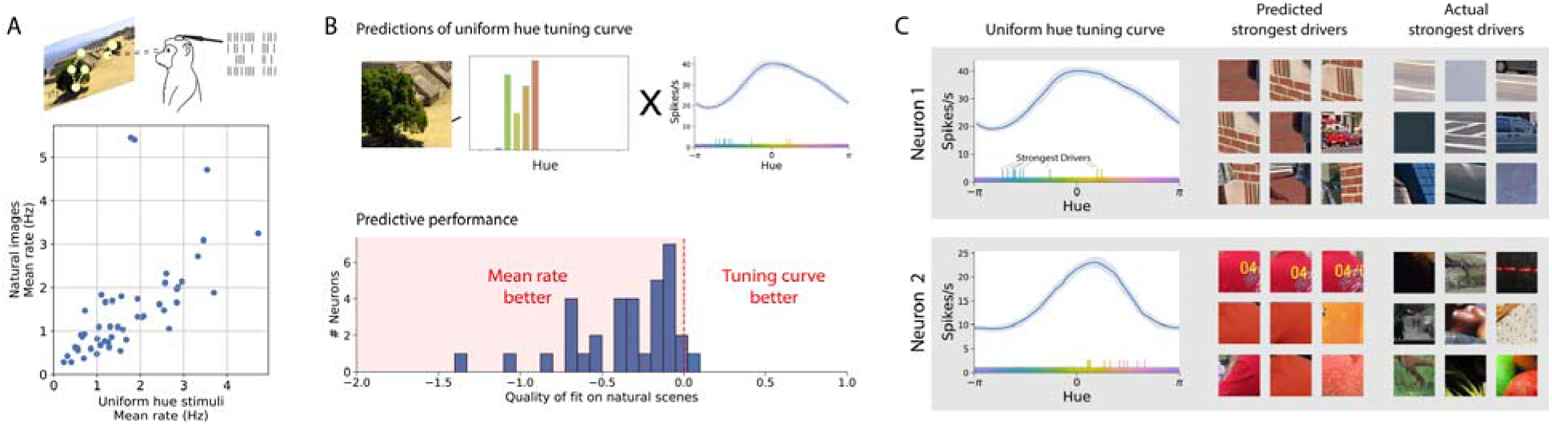
**A**) We displayed a large set of natural images to the same monkeys in interspersed sessions. Neurons fired at similar rates in these sessions as during presentations of a single hue. B) The hue-tuning on uniform hues (Fig. 1) can be treated as the coefficients of a linear model to predict neural responses to natural scenes, and scored. hue tuning on the artificial hue stimuli could not predict any variance in the natural scene responses. This is Displayed here is the histogram of the Poisson pseudo-R^2^ goodness-of-fit scores of the tuning curves’ predictions, which is below zero when the predictions underperform the mean firing rate. C) The tuning curves’ low predictive utility is exemplified in the disparity between the stimuli that they predicted strong responses for (e.g. the most red), and the stimuli that actually elicited the strongest responses, which were consistently of different hues.

### Natural scene models

We fit several models of the V4 response to natural scenes. Each of the models described below differs in the form of the encoding and the manner by which hue tuning curves are reconstructed.

#### Hue models

Our first model describes neural activity as a function of the hues present in the receptive field on each fixation. We used the same extraction of hues as above: we calculated the hue of pixels in the receptive field in CIELUV and binned these hues into histograms with 16 bins of hues. Since the hue histograms have 16 bins, the base regression problem to describe neural activity from hue is 16-dimensional.

As additional controls we included as covariates a small number of features unrelated to the images. To account for possible stimulus adaption, we included the trial number in the session and also the number of times the monkey previously fixated on that image. While all models predict the spike rate, which is already normalized by the fixation duration, we included the fixation duration as an input to control for possible nonlinearities of rate with fixation duration. We also included the duration of the saccade previous to the current fixation, the duration of the saccade after fixation, the location of the fixation, the maximum displacement of the gaze position during the entire duration of the fixation, and whether the pupil tracking was lost (often due to a blink) in the saccade before or after fixation. Including these inputs allowed the nonlinear methods to control for factors which also may affect spike rate.

For our nonlinear model, we selected the machine learning method of gradient boosted decision trees as implemented by XGBoost, an open-source Python package [41]. This method allows a Poisson loss function and has previously been shown to be effective in describing neural responses [42]. Briefly, XGBoost trains multiple decision trees in sequence, with each trained on the errors of the previous trees. We chose several regularization parameters using Bayesian optimization for a single neuron. These parameters included the number of trees to train (200), the maximum depth of each decision tree (3), the data subsampling ratio (0.5), the minimum gain (0.3), and the learning rate (0.08). The generalized linear model (GLM) presented in Supp. Fig. 2 (DOI 10.6084/m9.figshare.17957957) was a linear-nonlinear model with an exponential link function and a Poisson loss. We included elastic net regularization, and selected the regularization coefficient for each neuron using cross-validation with k=8 folds in an inner loop in the outer cross-validation for model scoring (see Model Scoring and Cross-Validation). We implemented this with the R package r-glmnet [43].

To build tuning curves from hue model we predicted the response to a vector indicating which color was present (that is, a “one-hot” vector with one entry per bin of hues that is all zeros except for the hue that is present). Then, to estimate the measurement error of the tuning curves, we refit the models to the original neural responses resampled with replacement (see Calculation of Error Bound). This resulted in tuning curves from hundreds of bootstrapped model fits. In figures in which we display the tuning curves, the lower and upper error bounds represent the 5^th^ and 95^th^ percentiles of the tuning curves observed when refitting the models to the resampled data.

#### CNN model

Our convolutional neural network (CNN) encoding model was based on findings that the intermediate layers of pretrained networks are highly predictive of V4 responses [5]. Ours was built from the VGG16 network, which is a large convolutional network trained to classify the images from the ImageNet dataset [44]. It contains 13 convolutional layers and 3 fully connected layers. We built an encoding model for each neuron from the activations of layer 14 (the first fully-connected layer), which we found to have the highest predictive power in conjunction with a nonlinear readout. We did not modify or refit this CNN to predict neural responses. Instead, we ran nonlinear Poisson regression (XGBoost) to predict each neuron’s response to an image from the values of layer 14 when the VGG network was given the same image. The final model thus takes a fixation image as input, runs the image through 14 layers of the VGG16 CNN, and then through a trained instance of XGBoost to predict the spike rate of a neuron. We call the combination of the CNN model and the trained XGBoost for each neuron the “CNN model”.

The CNN model could then be used to build tuning curves. We conceptualized this as extracting the average first-order effect of hue upon the responses of this model to natural images. We perform the following cross-validated procedure for each of 8 bins of hues. First, we train the CNN model (i.e. train the XGBoost regressor) on the training set of the natural image dataset. We then modify the test set images by slightly desaturating all pixels whose hue lies within the current hue bin. The bins were chosen to be large (8 in instead of 16) to so as to be less affected by pixel noise and to speed computation. We desaturated by multiplicatively reducing the chroma of colors in LUV color space, the same color space in which we define hue, by a certain factor. For robustness, we modified images at each of many desaturation levels, ranging from 5% to 100% of their original chroma. We then obtained the predictions of the CNN model to the original test set and also for each modified, desaturated test set, and take the average difference of these two predictions across all images. This process is repeated in an 8-fold cross-validation procedure, so that each image serves as the test set once. The resulting series of average differences can be plotted against the desaturation. The slope of this line represents the average first-order contribution of that bin of hues to the images in the dataset. Note that the value of slope reflects the scale the x-axis, which represents the parameterization of the desaturation percentage. It is best to think of the units of slope as arbitrary; the important result is the relative value of the slope between hues. Finally, the process was repeated for each bin of hues, resulting in the tuning curve to hue.

We sought to validate this procedure on simulated data (Supp. Fig. 4, DOI 10.6084/m9.figshare.17958104). One important aspect is that predictions are made on images that are as close to the distribution of images in the training set as possible. Since images in which a single bin of hues are desaturated by 5% are visually indistinguishable from the originals, this is not likely to be a concern. Nevertheless, we observed whether this method would be able to reconstruct the hue tuning of simulated neurons. We constructed 20 simulated neurons that responded linearly to the hues present in a receptive field. Each neuron was cosine tuned with a randomly selected hue angle. Linear regression could perfectly reconstruct the hue tuning of these simulated neurons, as expected. The CNN method could also reconstruct the tuning curves, though less well than linear regression. If linear tuning curves do exist, then, the CNN method would be able to reconstruct them.

### Receptive field estimation

To estimate hue tuning on natural scenes with the hue models, we needed to know which hues were present within the RF on each fixation. We mapped the RFs by presenting sinusoidal gratings at four orientations, which were flashed sequentially at the vertices of a lattice covering a portion of the visual field suggested by anatomical location of the implant. For monkey 1, this procedure identified an average RF over neurons in the implant of 5.87° in diameter (full-width at half-maximum) centered 8.94° below and 4.99° to the right of fixation, whereas for M2 we found an average RF 7.02° wide centered 7.02° below and 7.02° to the left of fixation. The location of the RFs were confirmed in the natural scene presentations as the pixel block that allowed the best predictions on held-out trials. For the hue model analyses, on each fixation we obtained the model inputs by extracting the hues present in the 50×50 pixel block (2.93° of visual angle on a side) surrounding the centroid of the RFs of each monkey.

We did not use this RF information in the CNN model, which took as input the entire image region around the fixation. Since information about spatial location preserved in the lower and intermediate layers of the CNN, the RF for any neuron can be learned. This addressed any worry that our conclusions are dependent upon the RF specification in the two hue models, and as well that the RF specification might systematically change for natural images.

#### Model scoring and cross validation

We quantified how well the regression methods described neural responses by calculating the pseudo-R^2^ score. This scoring function is applicable to Poisson processes, unlike a standard R^2^ score [45]. The pseudo-R^2^ was calculated in terms of the log likelihood of the true neural activity *L*(*y*), the log likelihood of the predicted output *L*(*ŷ*), and the log likelihood of the data under the mean firing rate *L*(*ӯ*).

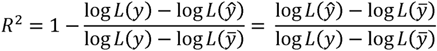

The pseudo-R^2^ is, at left, one minus the ratio of the deviance of the tested model to the deviance of the null model. It can be also be seen, at right, as the fraction of the maximum potential log-likelihood. It takes a value of 0 when the data is as likely under the tested model as the mean rate, and a value of 1 when the tested model perfectly describes the data.

We used 8-fold cross-validation (CV) when assigning a final score to the models. The input and spike data were segmented randomly by fixation into eight equal partitions. The methods were trained on seven partitions and tested on the eighth, and this was repeated until all segments served as the test partition once. We report the mean of the eight scores. If the monkey fixated on a single image more than once, all fixations were placed into the same partition. This ensures that the test set contains only images that were not used to train the model.

#### Calculation of error bounds

Each estimate of a tuning curve represents, in essence, a summary statistic of noisy data. To estimate error bounds on tuning curves, we relied on the nonparametric method of bootstrapping across trials, or for summary statistics of the entire neural population, additionally bootstrapping across neurons. Since the uniform field hue tuning curves used for correlations were simple averages of spike rates, binned over hue, we bootstrapped across trials to compute the confidence intervals. The natural scene tuning curves for the nonlinear hue model represented the predicted response to single hues. For these methods, we computed uncertainty bounds on their predictions to single hues by retraining the methods on resampled datasets (with replacement) and selecting the 5^th^ and 95^th^ percentiles of the predicted output for each bin. For the CNN method, the tuning curves were calculated from linear fits of the difference in test set predictions as a function of hue bin desaturation. The difference in predictions was noisy across images, with large changes predicted for some images but small changes predicted for other images. This noise presented as uncertainty in the linear fit to the data. The error on the CNN tuning curve, then, represented the uncertainty in the linear fit to the test set predictions.

The uncertainty on each of the tuning curves was then propagated into the correlation between the natural scene and uniform field tuning curves. This was again done through bootstrapping. For a given natural scene/uniform field correlation, we correlated the natural scene and uniform field tuning curves from hundreds of model fits upon resampled data, yielding a large distribution of correlations. We then reported the mean, 5^th^, and 95^th^ percentiles of this distribution. The uncertainty of the mean across neurons included a bootstrap across the trials used to build the tuning curves for each neuron, followed by a bootstrap across neurons.

#### Normative analysis of mixed selectivity

When neurons are nonlinearly selective for mixtures of a feature with others (a situation leading to tuning changing with context) they are said to have nonlinear mixed selectivity. Nonlinear mixed selectivity has previously been argued to be advantageous because it increases the dimensionality of the space of possible neural responses, which allows a greater diversity of linear readouts for downstream tasks [46]. Here we look to the optimal coding literature to find an alternative, more general justification. Our findings are summarized in Results.

A well-studied notion of optimality is that of Fisher efficient coding. In this framework the neural code is optimized to increase the Fisher Information it contains about the features of stimuli that are important for behavior. Denoting these features as ***θ*** = {*θ*_1_, *θ*_2_,…, *θ_M_*}, and the population activity of N neurons ***x*** = {*x*_1_, *x*_2_,…, *x_N_*} the Fisher Information is a matrix defined elementwise as:

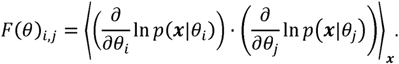

Here 〈·〉*_x_* denotes the expectation over given sources of noise. The Fisher Information is intuitively similar to the sensitivity of a representation across all neurons to a given dimension and value of ***θ***. We will ask what representations maximize the Fisher Information about all M encoded features.

Maximizing Fisher Information is a good measure of optimality because it describes how well any optimized decoder can read out the features ***θ*** from the response ***x***. Note that this normative reason is the *potential quality* of the readout, rather than the *overall number* of potential linear readouts as in the work of Fusi et al. [46]. Following the literature on optimal coding of neural populations [47–49], the decoding error can be bounded with the Cramer-Rao inequality [47]:

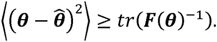

This states that the reconstruction error is lower-bounded by the trace of the inverse of the Fisher Information of the neural population with respect to ***θ***. By this metric, larger Fisher Information matrices allow lower error. Our analysis focuses on the case of neurons that fire with a mean rate *f*(***θ***) with additional independent Poisson noise. In this case the Fisher Information is a sum over neurons [50]:

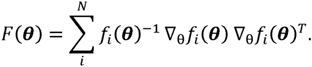

To show that mixed selectivity is advantageous in this setting, we will show that Fisher Information is larger (as measured by the trace) when neurons respond to mixtures of features rather than code for only one feature. Our approach rests on the fact that neurons with Poisson noise have lower variance at lower firing rates. Additionally, we assume that the features that neurons code for are sparse and often not present. We find that mixed selectivity is better in this setting because many neurons can participate in coding, rather than waiting in silence for their single feature, and because distributed coding allow lower firing rates for the same sensitivity.

Let the spike rates for the aligned response (one feature per neuron) be denoted as *f*(***θ***). A random rotation of this response is *Rf*(***θ***), where *R* is a random rotation matrix. We can additionally restrict R to those rotations that preserve positivity of the resultant spike rates. If neurons spike with independent Poisson noise, the Fisher Information of this mixture is

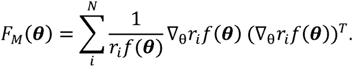

Here *r_i_* is the corresponding row of the mixing matrix. In this case the trace of the Fisher Information (our measure of coding quality) is:

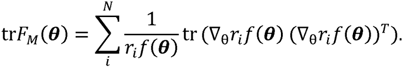

To ask if the rotation improves coding, we can ask if either of the two terms in the summand increase or decrease, on average. First, we find that the average of the right term does not change. This can be seen via the linearity of the trace and derivative and the cyclic property of the trace, 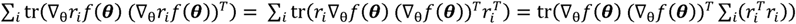. Since R is a rotation, *RR^T^* = *I*, and the average reduces to its value absent a rotation, 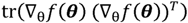. Thus, only the average of the scalar term due to Poisson noise 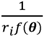 is affected by the rotation.

The value of 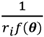 depends on the sparsity of the response. When neurons code for only one feature which is usually not present in a visual scene, most neurons are usually silent. The Fisher Information is controlled by the L active neurons. We can approximate the sum with 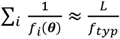 where *f_typ_* is the typical firing rate of the active neurons. This will increase for any random rotation with a probability that increases with the number of neurons. In fact, the dot product of a random vector such as *r_i_* with a fixed vector *f*(***θ***) concentrates around 0 in high dimensions, though this is complicated by the positivity constraint on *Rf*(***θ***). Taking a conservative approximation that the average firing rate of the rotated population is the average firing rate of the unrotated population, 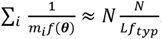. This is much larger than 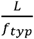 since *L* < *N*; specifically, a factor of 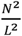 larger.

Thus, the Fisher Information is larger when the response utilizes all Poisson neurons at small firing rates, as in the rotated response, than when the response is concentrated on a sparse subset, as when each neuron codes for one feature. This makes coding for single visual features a disadvantageous strategy.

## Results

We recorded the spike rates of neurons in area V4 of two macaques as they viewed images on a monitor. One monkey (M1) freely viewed images as we tracked its gaze, while the gaze of the second monkey (M2) was fixed at image center during image presentation. We analyzed the responses of 90 neurons in M1 over several viewing sessions, taking care that the identity of cells on each electrode did not drift across sessions (see Methods: Session Concatenation), and in M2 recorded from 80 neurons in a single session. We then estimated tuning curves from responses to both artificial and naturalistic stimuli in order to ask if and how hue tuning generalizes.

### Tuning to hue on uniform screens

We first measured hue tuning by varying the hue of a uniform flat screen (Fig. 1A). We found that most of our neurons were well-tuned to specific hues, consistent with the previous literature on hue tuning in V4 [29, 30, 32]. Neurons’ strong selectivity for hues evenly tiled the hue circle (Fig. 1C). We characterized the degree of modulation with hue with the Modulation Index, calculated as the peak-to-peak range of the mean-normalized tuning curve (Fig. 1D). We also characterized our ability to estimate tuning by correlating the two tuning curves estimated on each half of the trials, selected randomly and bootstrapped for confidence bounds. The confidence interval of this correlation was usually high and excluded zero for 79/90 of neurons in M1 (Fig. 1C), but only for 17/80 neurons in M2 (Fig. 5A). A general trend across analyses was that neurons in M2 were more poorly described by hue than the neurons in M1. This difference in monkeys was possibly due to the spatial heterogeneity of color responses in V4 [29, 30]. In later analyses, we compared the hue tuning of neurons only when we could reliably estimate tuning. Thus, the placement of our electrodes in M1 and M2 identified neurons that, as characterized by stimuli of a single hue, appeared to robustly encode the hue of stimuli.

**Figure 1:**
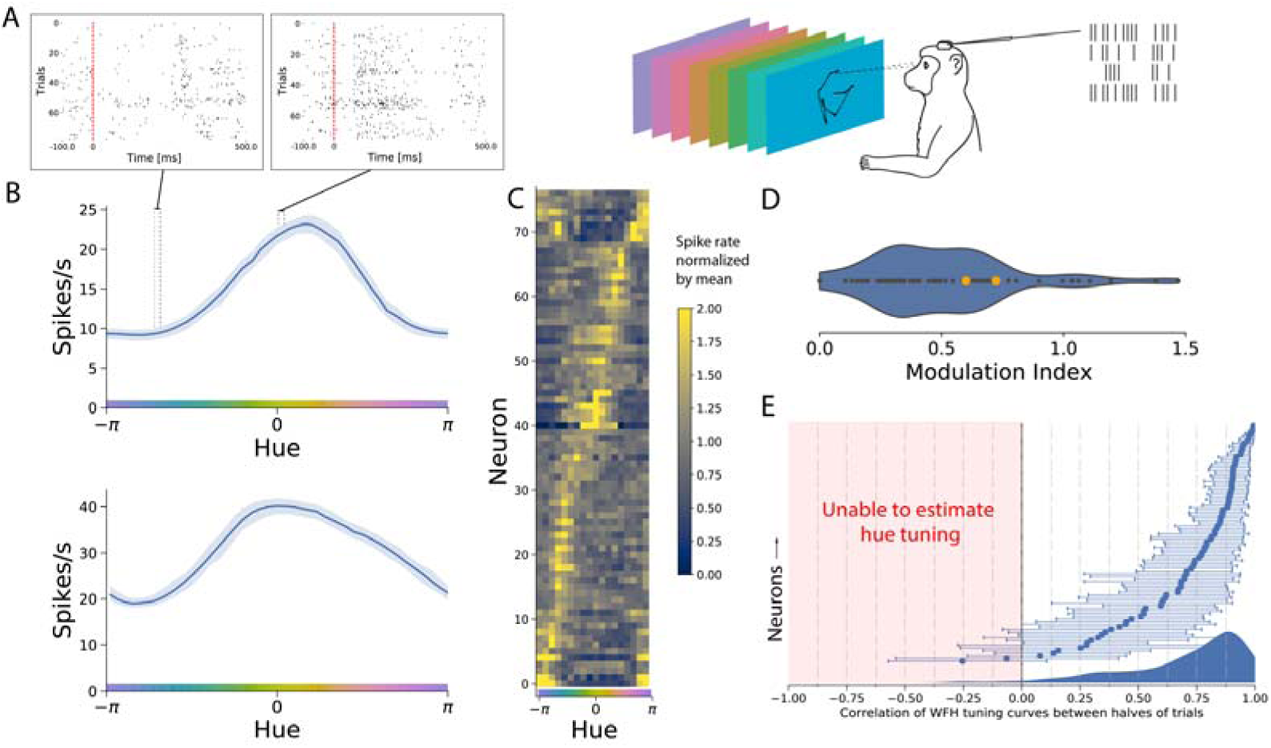
Tuning curves estimated from responses to artificial stimuli. Data from M1; see Fig. 5 for M2. A) We recorded from neurons in area V4 as a monkey viewed fields of a uniform hue and examined the average evoked spike rate 50-300ms after presentation. B) The uniform hue tuning curves for two example neurons, showing strong hue modulation, here displayed with LOWESS smoothing of trial responses. C) Most neurons modulated their activity strongly with hue. Here the unsmoothed tuning curves (mean rate in each bin of hues) are displayed normalized by per-neuron mean firing rate for comparison. D) The degree of hue tuning can be characterized with a Modulation Index, which is the difference in min and max of the tuning curve after normalization. The two example neurons of panel B are marked in orange. E) Our ability to reliably estimate hue tuning was captured by correlating the tuning curve estimated on one half of the trials with the tuning curve estimated on the other half. This correlation would be 1 in the no-noise or infinite-data condition, and if the 95% confidence bounds from bootstrapping include zero we cannot reliably estimate tuning.

**Figure 5:**
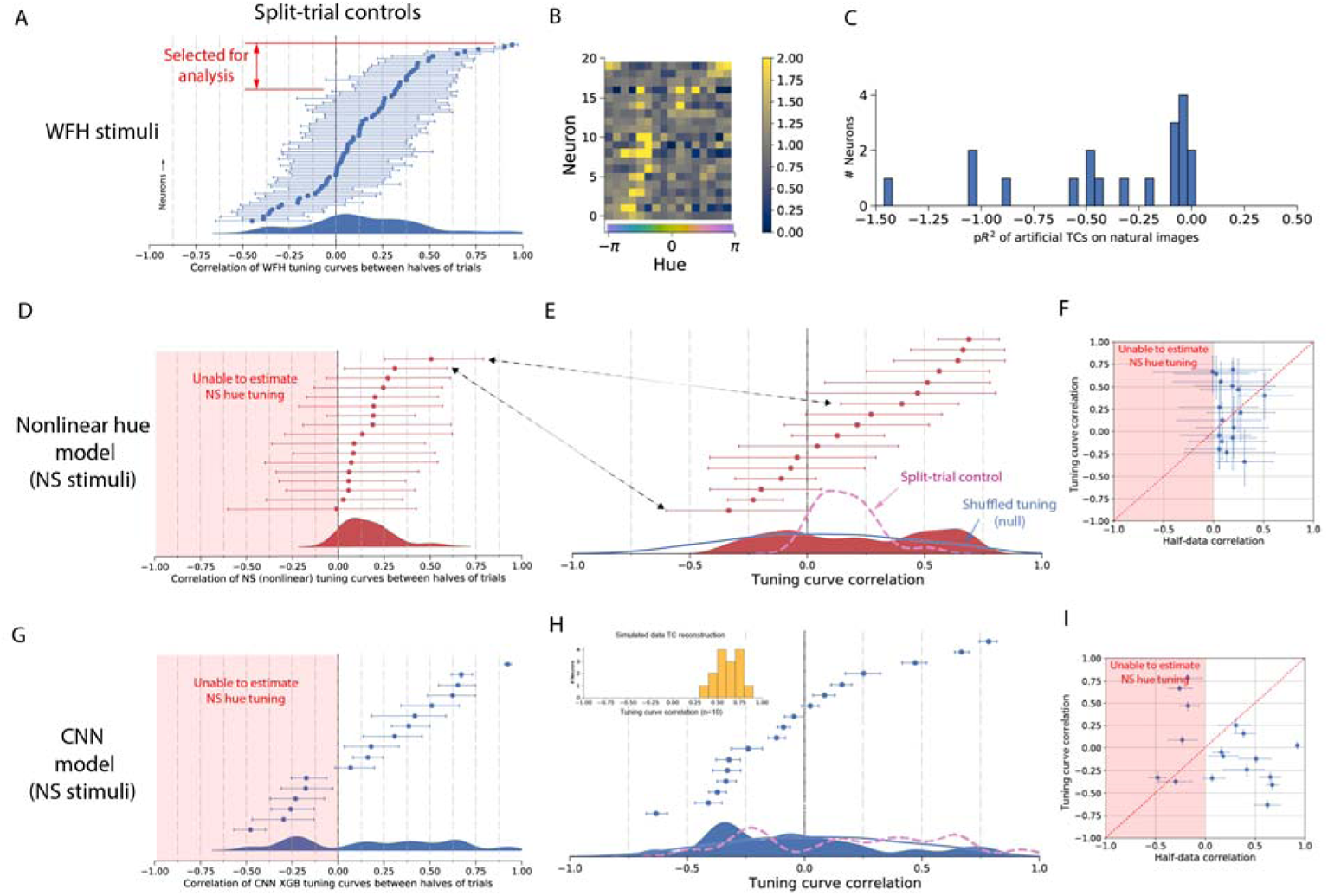
Collected data for M2. A) Most neurons in M2 showed poor hue tuning, and we were not able to consistently estimate uniform hue tuning nearly as well as for M1. B) Binned tuning curves for the neurons selected in A. C) As for M1, the uniform hue tuning curves were worse at predicting natural scene responses than the mean firing rate on natural scenes. D-F) Analysis of the natural scene tuning curves estimated by the nonlinear hue model was inconclusive. D) The natural scene tuning curves could not be estimated as consistently as for M1. E) The natural scene/uniform hue tuning curve correlations as estimated by the nonlinear hue model. Like for M1 (Fig. 2c) we overlay the split-trial distribution and the null distribution expected with random reshuffling of hue tuning. F) By a Wilcoxon signed rank test, we were unable to reject the null hypothesis that natural scene/uniform hue correlations are lower than the split-trial correlations (p=0.65) and thus it was not clear from the hue model on M2 neurons whether hue tuning does or does not change. G-I) Analysis of the natural scene tuning curves estimated with the CNN method. G) Distribution of correlations of hue tuning estimated of non-overlapping halves of trials. H) Natural scene/uniform hue correlations. Inserted is the distribution of natural scene/uniform hue correlations of simulated neurons with cosine hue tuning. Since M2 saw 10x fewer trials than M1, we simulated again on this smaller dataset. I) Among the neurons for which we could consistently estimate hue tuning (i.e. with a positive correlation of tuning curves estimated on split data), all neurons had a higher split-trial natural scene curve correlation than a natural scene/uniform hue correlation. This was significant under a Wilcoxon signed rank test at p=0.003.

If these tuning curves capture how these neurons encode hue, these tuning curves should predict responses to other types of stimuli. For example, we might expect that if a neuron preferred uniform fields of orange hue, then that neuron would on average have higher firing rates for scenes containing predominantly orange hues. To test this, we displayed natural images in alternating sessions to the same monkeys (Fig. 2A). We found that the tuning curves were not at all informative of the natural image response. Specifically, we asked how well uniform hue tuning curves could predict natural scene responses by interpreting the curves as the coefficients of a linear response to hue, and then scoring this model (see Methods). Our scoring method is a pseudo-R^2^ score which behaves roughly like an R^2^ but is valid on data with Poisson noise. Scores of 1 indicate perfect prediction, scores of 0 indicate the mean firing rate is an equally good predictor, and negative scores indicate the mean rate is a better predictor of firing than the model. Of all but one neuron, the uniform field tuning curves predict natural responses with a negative pseudo-R^2^ (Fig. 2B). Thus, neurons tuned for a certain color presented in isolation did not on average fire more when that color was present in natural scenes.

These observations can be explained if the hue tuning curves themselves are different between two contexts. Alternatively these neurons could respond more strongly to non-hue features that co-vary within natural images, like the visual texture of typically green plants. To distinguish these two possibilities, we next estimated tuning to hue directly from the responses to natural images and compared it with uniform hue tuning.

### Tuning to hue estimated from natural scenes

To investigate if hue tuning changes in the context of natural images, we directly regressed the contribution of hue to the neural response using two separate models. These models are of varying complexity and nonlinearity. Collectively they control for other visual features that drive V4 neurons, including interactions between hues, and rule out the possibility that visual confounds could explain the discrepancy between uniform field hue tuning and natural scene hue tuning.

#### Nonlinear model with hue as an input feature

We first modeled natural image responses as a nonlinear function of the hues present in the receptive field of neurons during each fixation (Fig. 3A) plus control features to account for effects such as adaptation. This model, which we refer to as the ‘nonlinear hue model’, predicted neural activity during natural scenes quite accurately for neurons in both monkeys (Supp. Fig. 2B,C; DOI 10.6084/m9.figshare.17957957). We fit a nonlinear model because nonlinear hue interactions have been previously observed in V4 [51], which would lead to a bias in a generalized linear model (GLM) because hues are correlated in natural scenes (Supp. Fig. 2A). Indeed, the nonlinear model was much more accurate than a GLM fit to predict spike rates using hues of natural images (Supp. Fig. 2B,C). This approach to estimating hue tuning directly regresses the response to bins of hues in natural images.

**Figure 3.**
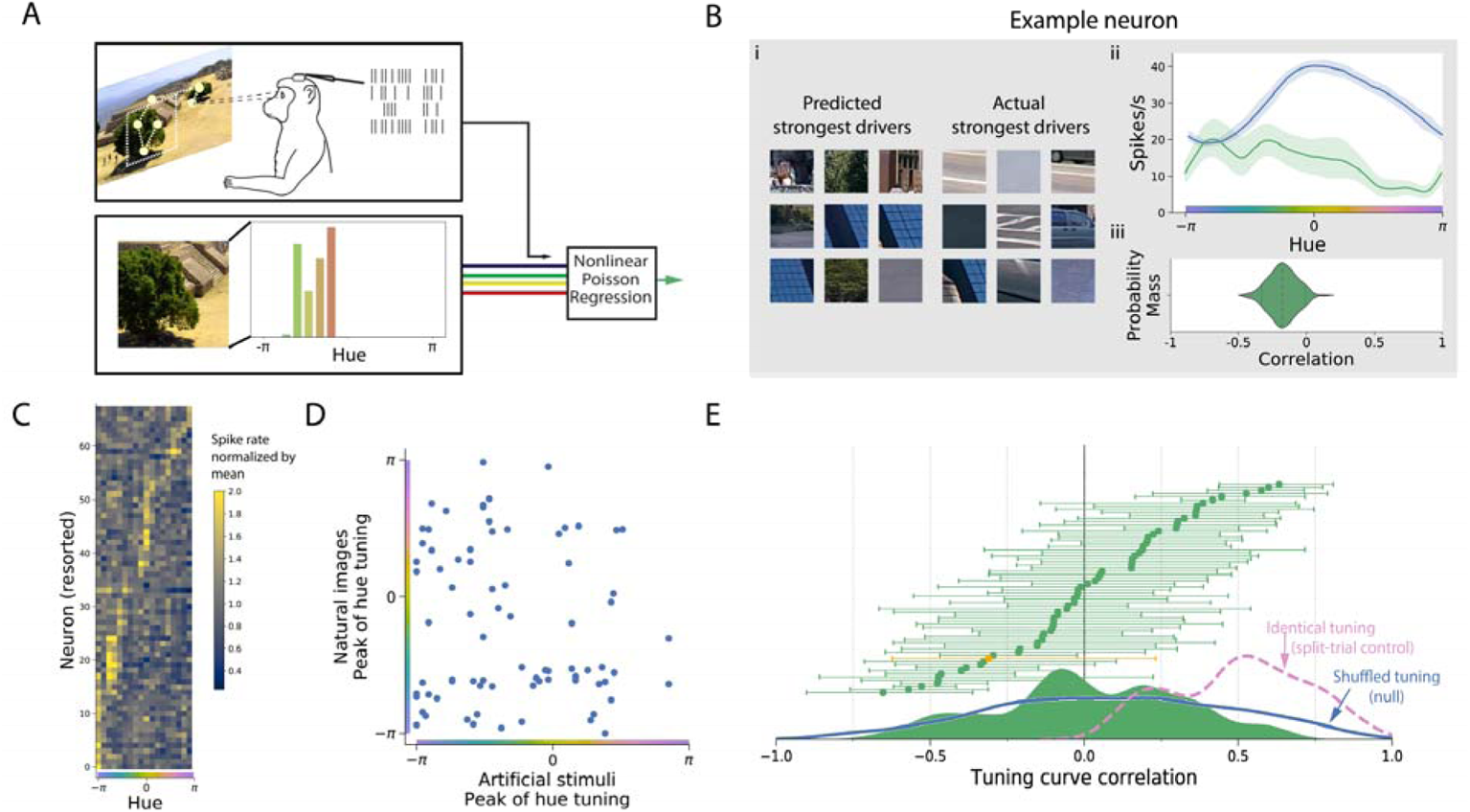
Tuning curves for hue estimated from the responses to natural images. Data from M1; see Fig. 5 for M2. A) We trained a nonlinear model with Poisson output to predict each neuron’s response from the hues present in its receptive field during the natural scene sessions. B) (i) The 9 trials that each model predicted to have highest firing rate looked similar to trials with the actual strongest response, unlike the uniform hue model. (ii) We built tuning curves from the model by observing its response to a single hue. The uncertainty of each curve is given by the 5^th^ and 95^th^ percentiles of hundreds of model fits to the trials resampled with replacement. (iii) This uncertainty is then propagated into the correlation between the uniform hue tuning curves and natural scene tuning curves. C) Tuning curves across all neurons (resorted by hue tuning in this condition). D) The peaks of these tuning curves plotted against the peaks of the tuning curves in the uniform hue condition show no circular correlation across neurons, *p*=0.82. E) The correlations of the natural scene and the uniform hue tuning curves on each neuron (as shown in B(ii) and (iii)) show this is not an artifact due to multiple peaks in tuning curves. The neurons are sorted by their correlation to show a cumulative distribution. The two example neurons are highlighted in orange. Below: The smoothed density of all neurons’ natural scene/uniform hue correlations is similar to what would be expected if neurons randomly shuffled hue tuning between conditions (overlaid, blue). Also overlaid (in pink) is the control distribution of how the correlations might appear if tuning were the same across stimuli, which is limited by neural noise and finite trials. This is estimated conservatively by correlating tuning estimated from one half of the natural scene trials with the tuning estimated on the other half.

We estimated hue tuning curves for the nonlinear hue by measuring its responses to single hues, in essence reproducing the uniform hue experiment but on the natural scene model. These tuning curves showed clear preference for small ranges of hues (Fig. 3C). We quantified our ability to estimate hue tuning in two ways. First, we repeatedly refit the model on the natural scene trials resampled with replacement, and observed the distribution of coefficients (Fig. 3Bii). This distribution was propagated through to later analyses such as the correlation between a neuron’s hue tuning estimated in either stimuli set. Secondly, we visualized how high the correlation of hue tuning across conditions would have appeared if tuning were the same in both contexts, given all sources of noise and measurement error. This we estimated by comparing the hue tuning curves from two non-overlapping halves of natural scene trials (Supp. Fig. 1; DOI 10.6084/m9.figshare.17957408). Note that this split-trial control is a conservative lower bound of our quality of estimation, as the model was fit on only half the number of trials. By this measure, the nonlinear hue model was able to consistently estimate hue tuning for the most neurons in M1 (Supp. Figure 1A, see also Fig. 3E) but for just two neurons in M2 (Fig. 5 D-F), which prevented a statistical analysis in M2. These estimates of uncertainty serve as a baseline limit of how well we can observe changes in hue tuning.

We compared the tuning curves across stimulus sets for neurons for which we could consistently estimate hue tuning. The peaks of the tuning curves did not show any correlation across neurons (Fig. 3D; circular correlation of - 0.02, unable to reject the uncorrelated hypothesis with *p*=0.82). In addition, the shapes of the tuning curves did not correlate between conditions (Fig. 3E). If hue affected V4 responses in the same way in both contexts, we would have observed the correlations between tuning curves across contexts to be at least as positive as the split-trial control. This was not the case. In M1, the natural scene/uniform field tuning curve correlations were significantly lower than these split-trial correlations (p=1.0×10^−14^, Wilcoxon signed-rank test; Supp. Fig. 1D, DOI 10.6084/m9.figshare.17957408), indicating that the observed change in hue tuning across contexts was not a consequence of noise in the estimation of tuning. In fact, the spread of correlations between the two sets of tuning curves was similar to the distribution that would arise if hue tuning shifted randomly between contexts (Fig. 3E inset), which to preserve typical tuning shapes we approximated as the correlations between random neurons’ tuning. Thus, the regressed contribution of hue to the neural response had little relationship to the strong tuning observed in response to stimuli of a single hue.

#### Neural network model of V4 responses

We next repeated the estimation of hue tuning on natural scenes with a more general model of V4 neurons that does not rely on hand-specified summaries of the features present in a receptive field. This was important to ensure that our results were not sensitive to design decisions in processing the images, as well as to account for the confounds of other, non-hue features contained in the image. The two hue models would provide biased estimates of tuning if neurons also responded to other visual features, and if these features co-varied in the image dataset with hue. If most green objects are plants, for example, the observed dependence on green hues may be partially attributable to a response to the high spatial frequency of greenery. Theoretically, one could include these features as additional covariates, but the list of features that drive the V4 response in a simple manner (e.g. linearly) is not comprehensively known. Good progress has been made with shape and texture [3, 52, 53], but arguably not enough to thoroughly control for all non-hue features in a simple model. Controlling for other drivers of V4 thus requires a model that learns relevant visual features instead of using features chosen by a researcher or parameterized by hand.

The model we selected was based on an encoding model of V4 that relates neural responses to the activations to the upper layers of a convolutional neural network (CNN) pretrained to classify images [5]. Such “transfer learning” models have also recently been used to infer optimal stimuli for neurons in V4 [18, 54]. Instead of pre-specifying a receptive field estimated with sparse noise, we allowed the CNN model to learn any location sensitivity itself and thus fed the entire fixation-centered image as input. Predictions of neural activity are obtained by passing the image through the network, obtaining the intermediate network activations, and then passing these to a classifier trained for each neuron (Fig. 4A). The predictions of neural activity given by this model were comparable in accuracy to those of the nonlinear hue model (Fig. 4B) despite the model making many fewer assumptions about how raw pixels related to responses.

**Figure 4.**
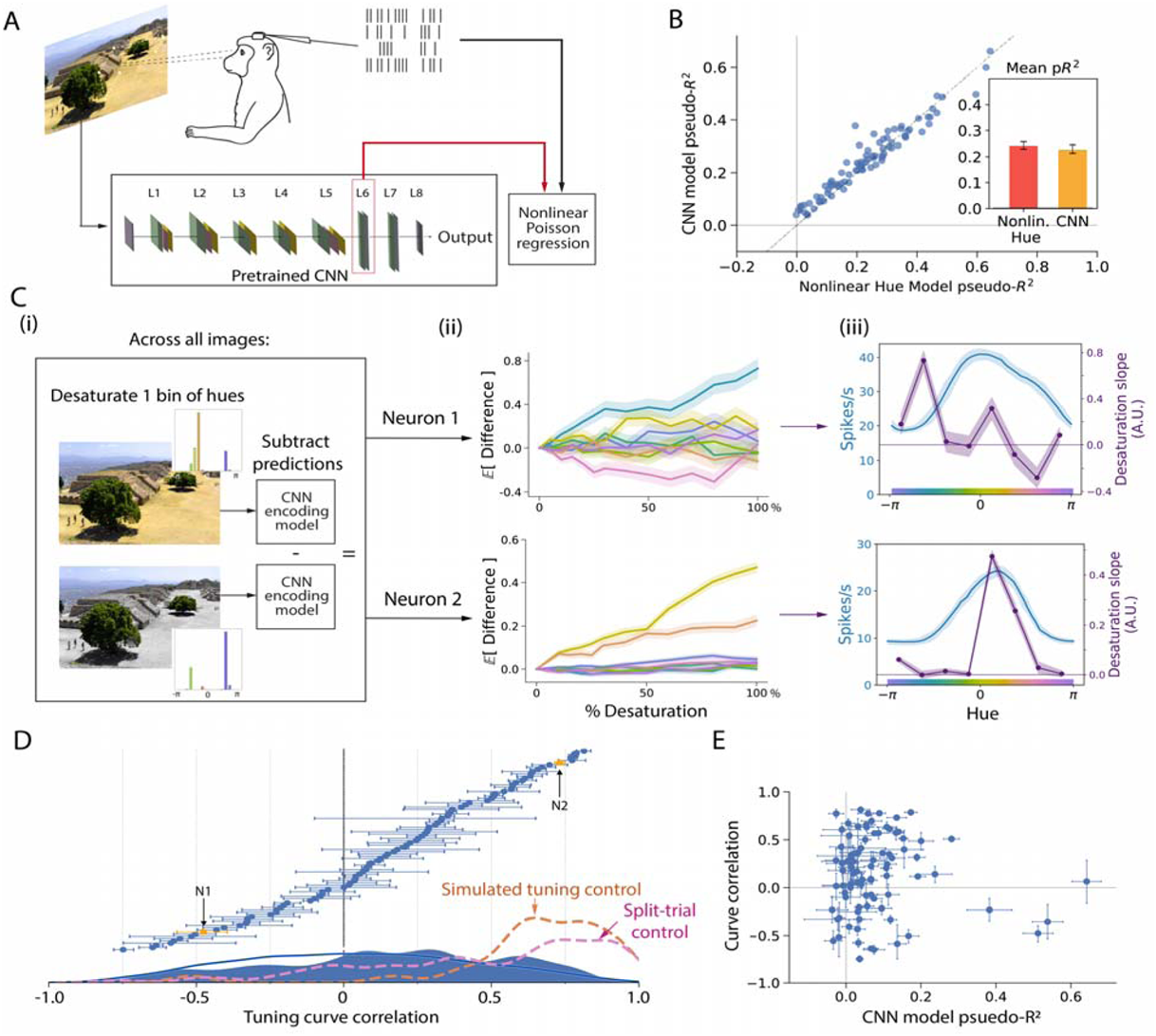
Tuning curves estimated for hue from a model of V4 responses built from a pretrained convolutional neural network (CNN). Data from M1; see Fig. 5 for M2. A) We trained a nonlinear Poisson regression model (gradient boosted trees) to predict the V4 response from the activations of an intermediate layer in the VGG16 network given the visual stimulus. B) The quality of the neural predictions on each neuron, measured by the cross-validated pseudo-R^2^ score, were similar between the CNN model and the nonlinear hue model. C) We built hue tuning curves in the following manner: (i) For each image in a test set, we slightly desaturated all pixels in a bin of hues, and subtracted the CNN model’s predictions on the perturbed image from those on the original image. (ii) For each neuron, the average change in the predicted response across all test images was plotted against the percentage by which hues were desaturated. The slope of each line is, to first order, the average effect of that hue on the model response in the test set. The top and bottom plots show the same example neurons as in earlier plots. (iii) The resulting tuning curve (purple) summarizes the average effect of each of the 8 bins of hues – i.e. the slopes of the 8 desaturation curves. It can be seen that the tuning of neuron 1 was poorly correlated with the uniform hue tuning (blue), while that of neuron 2 was well-correlated, in agreement with the hues of the strongest-driving stimuli shown in Fig. 1B. D) We calculated the correlation between the two tuning curves for all neurons. The distribution of correlations was lower than for the reconstructed hue tuning of simulated neurons (“simulated tuning control”; see also Supp. Fig. 3) as well as the distribution of correlations between tuning curves estimated from two non-overlapping halves of the natural scene trials (“split-trial control”; see also Supp. Fig. 1). E) The quality of the CNN model fit for each neuron did not predict the correlation of the tuning curves.

Our initial, unsuccessful method to estimate hue tuning from this model was to simply observe the model’s response to images of a uniform hue, as before. However, this approach failed to reconstruct tuning on simulated data. This interesting parallel to our main finding is likely due to the fact that uniform field test images are far outside the domain of natural scenes on which the CNN was pretrained.

Instead, we developed a method to estimate hue tuning from the model that only uses responses to images close to the domain of natural images. By slightly perturbing the hue of input images and observing the change in the learned model’s response, we could test the model’s sensitivity to hues to in natural images (Fig. 4C). First, for a test set of images not used for training, we desaturated all pixels within a bin of hues by a set percentage (Fig. 4Ci). The percentage of desaturation varied from 0% (i.e. no change) to 100% (in which all pixels of one hue are taken to the isoluminant grey). We took the difference between the model’s predictions on the original and perturbed images and examined how severely this difference depended on the level of desaturation (Fig. 3Cii, iii). For each neuron, we averaged over the entire image dataset to yield the average effect of perturbing each hue on natural images. This method established the effect of hue only in the tight neighborhood of each image, and is set up to estimate the average local effect of hue on the natural image response.

To ensure that this process could in principle reconstruct correct tuning curves, we built simulated responses (Supp. Fig. 3; DOI 10.6084/m9.figshare.17958104). We generated random cosine tuning curves, then simulated a hue response by applying these as linear filters upon the histograms of the hues present in each image. We then attempted to predict these simulated responses from the activations of the pretrained CNN given the raw images. Using the method of progressively desaturating test images, we found we could reconstruct the original cosine tuning curves with high accuracy (Fig. 4D overlay and Supp. Fig. 3). As a second, more conservative test, we also performed the split-trial control for the actual V4 neurons, which involved repeating the entire analysis separately on two non-overlapping halves of natural scene trials and then correlating the two resulting tuning curves. The split-trial tuning curves showed significantly positive correlations for most neurons in M1 (Fig. 4D overlay) and M2 (Fig. 5). This method of querying the effect of hue could thus accurately estimate hue tuning curves from natural scene responses in both monkeys.

We next asked if these tuning curves would be similar to the tuning curves to uniform hues. We found that the tuning curves of one context were different from tuning in the other (Fig. 4D for M1 and Fig. 5G for M2). Among those neurons for which we could consistently estimate hue tuning, the natural scene/ uniform hue tuning curve correlations were significantly closer to 0 (p=1.1×10^−8^, Wilcoxon signed-rank test, Supp. Fig. 1D for M1; Fig. 2H for M2). This difference in tuning curves was not an artifact of our model fit or estimation method, as this would be measured in the split-trial control, and additionally we observed no correlation between the model’s accuracy on unseen natural images and the natural scene/uniform field correlation (Fig. 4E and Fig. 5I).

In addition to changes in tuning curve shape as captured by correlation, we also examined if the natural scene tuning curves showed changes in the overall degree of hue modulation. We found that hue modulation – the maximum of a tuning curve minus the minimum, normalized by the mean – was related across contexts, but weakly (Supp. Fig. 4; DOI 10.6084/m9.figshare.17958221). Many neurons strongly modulated by hue on uniform fields had weak responses to hue on natural scenes, and vice versa. Overall, the tuning curves estimated with this more advanced method support our previous conclusion that hue tuning on uniform fields does not agree with the effect of hue in natural scenes.

### Why features interact

A straightforward explanation of why hue tuning differs across visual contexts is that these neurons respond to nonlinear combinations between hue and non-hue features, as shown schematically in Figure 6. What computational advantage could explain this coding scheme for visual perception? It is clear that if the role of these neurons were to encode hue alone, then any nonlinear interactions would be detrimental. This is because hue can no longer be unambiguously read out without additional contextual information. Therefore these V4 neurons likely assist in a more general task, like object recognition or segmentation. Other studies have also noted that color vision may be best thought of in terms of task performance [55]; the absorbance spectra of the L and M photoreceptors in primates, for example, are not maximally separated as in birds but rather overlap significantly, possibly because this helps to discriminate and classify fruit and leaves [56]. The question then arises: why would neurons being responsive to multiple features help visual processing?

**Figure 6:**
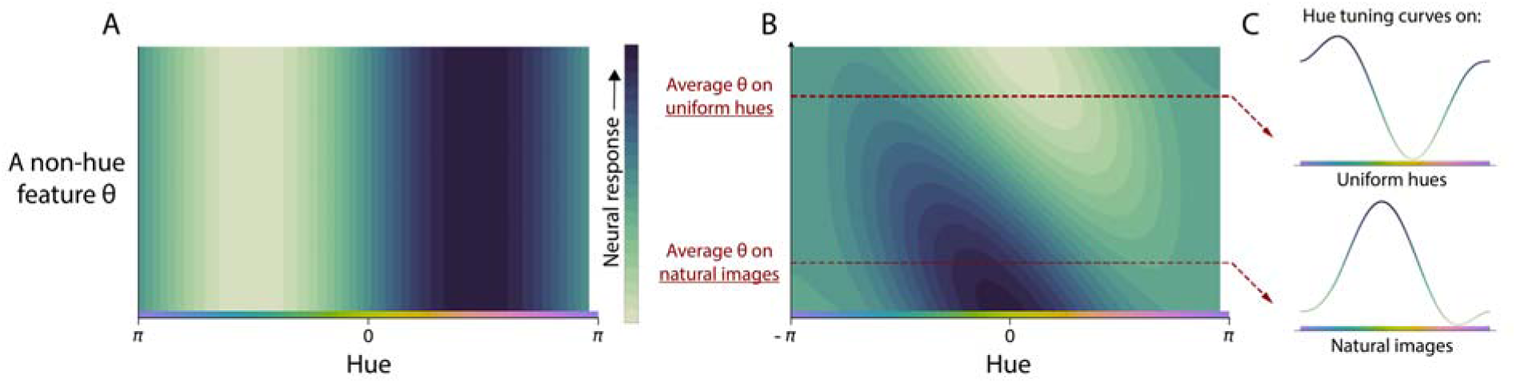
Interactions between features allow neurons to carry more information in their activity. A) In this two-dimensional tuning curve, a hypothetical neuron responds to only hue and carries no information about other variables. B) A hypothetical neuron that additionally responds another non-hue feature is informative about multiple dimensions of stimuli (due to its nonzero derivative). C) We can build a hue tuning curve for this neuron by varying hue with the other feature held fixed. If the average non-hue feature is different between natural images and uniform hues, the tuning curves to hue will differ between contexts.

One possible reason is that selectivity to multiple features increases the dimensionality of the space of possible neural responses, which allows a greater diversity of linear readouts for downstream tasks [46]. This justification prioritizes the diversity of possible uses of an area rather than the accuracy of encoding. Here, we look to the optimal coding literature (e.g. [47, 50, 57]) to find an alternative normative justification. Starting with a certain set of behaviorally-relevant visual features, what is the optimal way of representing these M features in a single population of N neurons?

Our findings, derived in Methods, show that mixed selectivity allows for better neural codes when neurons fire as Poisson processes and visual features are sparse and often not present. This is because a mixed selectivity strategy allows many neurons to participate in each response, rather than a sparse subset. When each neuron responds to *k* features, *k* times more neurons can respond on average to each scene. A distributed response in turn enables lower firing rates for the same sensitivity, which is advantageous because Poisson noise has lower variance at low firing rates. Coding quality, which is related both to the variance of internal noise and the sensitivity of the response when measured by Fisher Information, thus improves when the response is maximally distributed across many neurons. Recent studies using other measures of coding besides Fisher Information support this conclusion that mixed selectivity improves neural encoding [58].

It should be noted that an optimal coding argument may not necessarily explain selectivity to all features. Orientation selectivity, for example, is similar across contexts [11] and this may be related to fundamental visual cortical architecture and the importance of visual form for behavior. Where neurons do not show mixed selectivity, anatomical or behavioral constraints may override mixed selectivity’s benefits of increased precision and quality of the neural encoding.

## DISCUSSION

For populations of V4 neurons in two macaques, we found that varying the hue of simple stimuli produced tuning curves that do not accurately describe hue tuning measured from natural scenes. While some discrepancy may be expected, we found that the two sets of tuning curves correlated not much better than chance. This finding was robust across multiple methods of estimating tuning, which together accounted for the confounds of both hue-hue interactions as well as of non-hue drivers of V4 activity. A hue tuning curve for V4 estimated from any one set of stimuli thus does not universally describe the average response to hue on other stimuli.

### Known sources of modulation in visual responses

The V4 response is modulated by a number of factors that change with visual context. These factors are divided in the manner of their relevance to our findings. First are possible reasons why we might have observed low tuning curve correlations even if, in fact, tuning did not change between contexts. The second category of factors are known interactions between hue and other features in the V4 response that may explain why hue tuning in V4 changes with visual context. We will review both in turn.

Of first concern as a potential confound upon hue tuning estimation is visual attention [59]. A particularly relevant form of attention is feature-based attention, in which neurons tuned for a feature (say, red) increase their firing rate if that feature is attended to (as in the task, “find the red object”) [60, 61]. While the task of M1 was free viewing and involved no instructions, it is likely that the monkey’s attention shifted during the task and that it was influenced by object salience. This effect may bias our results if object salience were correlated with hue. Attention is less likely to have presented a confound in the task of M2, in which gaze was fixed at center and stimuli were presented for 100ms. We have not directly controlled for attention, apart from trends in salience that might have been learned by the CNN model, but we believe that the size of the apparent change in hue tuning cannot be attributable to salience-hue correlations.

Neurons in V4 are have been shown to preferentially respond to objects near the center of attention, even when attention falls away from fixation [62–64]. This phenomenon of receptive-field remapping is most problematic for our hue models, which required that we extract the hues lying within the receptive field. If the monkeys’ attention frequently strayed away from fixation, we would have extracted hues from an irrelevant image portion. This would introduce some noise in the hue covariates, and therefore some smoothing of hue tuning curves. The CNN model learned any spatial sensitivity directly from the natural scene responses. However, the effect of attention upon receptive fields could not be modeled and it is likely that some smoothing of the hue tuning curve occurred for this technique as well. Smoothing would obscure fine-scale structure in the tuning curves. As the curves were already smooth, however, the natural scene/uniform field correlations should not be much diminished. The smoothing effect is furthermore not consistent with our finding that many neurons have natural scene hue tuning with zero, or even negative correlation with their uniform field tuning while still showing strong hue-dependent modulation.

We now turn to potential descriptions of the interactions that might have led to a shift in hue tuning across contexts. One possibility is the behavioral phenomenon of color constancy, which would present as a neural correlate as responses to the inferred surface color of objects rather than their apparent color (which reflects the color of ambient light) [51]. This is a clear example of the nonseparability of the V4 response to hue, and a reason hue tuning might change between any two, single stimuli. It is less obvious, however, that color constancy correlates would cause the average effect of hue over all natural images to be different than on uniform hues. It would be expected that over tens of thousands of images with a broad range of lighting conditions, color constancy would result in some smoothing of the estimated tuning curve due to the difference between the pixels’ hue and the inferred hue, and of the same characteristic scale as their typical difference. Additionally we may expect a bias that would result from the discrepancy between pure white and the average lighting condition. We expect this discrepancy to be small, and therefore that natural scene tuning curves would still be strongly (though not perfectly) correlated with the uniform field tuning curves. Though phenomena like color constancy would affect hue tuning on natural scenes, it cannot account for the entire difference we observed, and it is likely that there exists other undocumented sources of nonseparability.

A subpopulation of neurons in V4, so-called equiluminance cells, respond to object boundaries defined solely by chromatic boundaries [65]. Such shapes are defined by changes in hue or saturation, and so it is worth asking whether the response function of equiluminance cells includes interactions between hue/saturation and spatial arrangement. However, it was not originally determined if the responses were actually separable in this way, as neurons’ hue tuning curves were characterized with a fixed shape. It is possible that equiluminant cells had fixed hue tuning that was then modulated by shape. Thus, it is plausible but undetermined that equiluminance cells would show different hue tuning across shape and explain our results.

The apparent shift in hue tuning in natural scenes may be partially be explained by a multiplicative or generally nonlinear interaction between shape and color, as was examined in a recent paper that jointly varied the hue and shape of simple stimuli [66]. In a linear model with terms for hue and a multiplicative interaction between hue and shape parameters, the authors observed a significant interaction between shape and color in the majority of cells (44/60). This interaction would cause (linear) hue tuning to appear different for natural images with varying shapes, as we observe. We note that other, undescribed features may also interact with hue, and that conclusively determining which visual features interact would require presenting stimuli tiling many more dimensions of variation.

### Implications for V4 and for the tuning curve approach

Color responsivity has long been a defining feature of V4 [19, 67]. Recent studies have shown that localized areas in V4 are strongly responsive to color [29], and furthermore that the anatomical organization of color preference on the neocortex is similar to perceptual color spaces [31–33]. These findings have been taken as evidence that areas within V4 are specialized for the perception of color. However, each of these studies characterized hue tuning by changing the color of simple shapes. Since the color tuning of V4 neurons changes with visual context, as we show here, it is possible that previous conclusions about the functional organization of V4 do not accurately describe how V4 processes more naturalistic stimuli.

It should be noted that our simple stimuli were not colored shapes chosen by hand to drive V4 neurons strongly, as in many previous studies, but rather uniform screens. We found that these still elicited strong responses and well-identified tuning curves. Nevertheless it may be objected that previous studies’ stimuli may still measure hue tuning that generalizes to natural scenes. However, this would require that the factors that modulate hue tuning in V4 neurons are only present in uniform screens. It is more parsimonious that hue-tuned neurons are modulated by interactions with a range of spatial features, which collectively will cause tuning on any set of stimuli to not generalize to naturalistic stimuli.

Based on the discovery of robust tuning for the color of simple visual stimuli, some studies have concluded that the role of color-responsive areas in V4 is to represent color. Our results do not rule this out; for example these areas might represent color but be modulated by what colors are likely given the surroundings. This would complicate a read-out of color from V4, but may have other advantages like efficiency. It would be interesting to investigate this possibility in future studies. An alternative possibility is that the color-sensitive areas of V4 are not specialized to represent color, *per se*, but rather serve a more complex role within recognition and perception. This is analogous to how V2 appears tuned to orientation but can perhaps be better described as processing naturalistic texture [68]. Furthermore, this role aligns with the suggestion that the ventral temporal cortex at large decomposes scenes into neural activity such that object categories are linearly separable [69]. Thus, the color-responsive areas of V4 may represent how color informs an inference of object identity [55]. Whether the color responses of V4 are an end to themselves (i.e. representing color) or intermediate computations in a larger assessment of object identity [70], or both, cannot be decided from this study; both are consistent with the data.

Our study joins a longer history of literature observing that, across many brain areas, tuning curves previously characterized with simple stimuli in fact change with context. In V1, for example, researchers found that receptive fields change with certain visual aspects that were not varied within previous stimuli sets, such as the presence of competing orientations [71–74]. Even sound has been shown to modulate V1 receptive fields, at least in mice [75]. More recently, it was observed that receptive fields are different in the contexts of dense versus sparse noise for neurons in layer 2/3 of V1 [76]. Spatio-temporal receptive fields of V1 neurons also appear different when estimated on natural movies versus drifting gratings [10, 12] (though note that orientation tuning is similar for static natural scenes versus gratings [11]). In other areas, such as for retinal ganglion cells [77–79] and in macaque M1, S1, and rat hippocampus [42], contextual modulation in the form of nonlinear feature interactions have been identified by showing perturbed natural images instead of white noise or by comparing the performance of a model that assumes separability (such as a GLM) with a nonlinear model that does not. Thus, while tuning curves generalize in some situations (e.g. [11]), it is common that they do not, and any assumption of separability of the neural response should be verified. Furthermore, as derived in Methods, feature interactions are likely optimal for visual processing when the full visual scene is represented in neural activity and should be expected. Unless specifically investigated, it might not be correct to assume that a tuning curve accurately describes the neural response on different stimuli than used to create it.

If it cannot be assumed that neural tuning is separable, however, it becomes necessary to test prohibitively many stimuli or else make an alternative simplifying assumption. This is because the stimuli must scatter the entire space of relevant features, rather than be systematically varied along just one feature at a time. Since the number of tested stimuli must follow the number of potential feature combinations, the overall number of stimuli will grow exponentially with the number of features. When there are very many features, even very large recording datasets by today’s standards may be insufficient.

One possible way forward is to make simplifying assumptions, i.e. to set strong priors of the kinds of tuning curves that could be expected. This is the approach taken, for example, when modeling neurons using the activations of deep neural networks pre-trained on image classification tasks [5, 80] or considering neural responses as implementing a sparse code [9, 81]. To compare with the previous literature, single dimension experiments can then be performed on these complex encoding models, as we demonstrate here, or alternatively performed directly on artificial neural networks to gain intuition about what tuning curves say about information processing [82, 83]. In general, finding suitable priors will require the use of strong theoretical ideas and mechanistic hypotheses. To estimate tuning without assuming separability, then, neurophysiology must embrace and develop theories of neural processing.

## Acknowledgements

We are grateful to Samantha Schmitt for assistance with data collection and spike sorting. K.K., P.R., H.F., and A.B. acknowledge support from NIH grants MH103910 and EY021579. M.S. was supported by NIH EY029250, MH118929, EB026593, and NSF NCS 1954107.

## Author Contributions

A.B. prepared the modeling methodology, conducted analyses, prepared figures, and wrote the initial manuscript. P.R. and H.F. conceptualized the research, designed the experimental protocol, curated the data, prepared methodology and initial analyses, and edited the manuscript. M.S. provided funding and resources for the experiment, designed the experimental protocol, performed the data collection, advised the analysis methodology, and edited the manuscript. K.K. conceptualized the project, provided funding, advised the experimental protocol, supervised the data analysis, and edited the manuscript.

## Data Availability

All electrophysiological data is available upon request. Data can be provided in various stages of preprocessing to expedite replication: as spike times in each session, or as spike counts paired with raw images, with fixation-centered images, or with extracted hue histograms.

## Code Availability

Our in-house code for spike sorting is available at https://github.com/smithlabvision/spikesort, and the code used to process sorted data, run analyses, and create figures is available at https://github.com/KordingLab/V4py. Our analyses depended on Python v2.7 and a large number of open-source projects that can be found listed in https://github.com/KordingLab/V4py/requirements.txt, including Pandas v0.23, Tensorflow v1.1, Numpy v1.11, and Keras v2.2.

