## Supplemental Figures for "Hue tuning curves in V4 change with visual context"

### Supplementary information

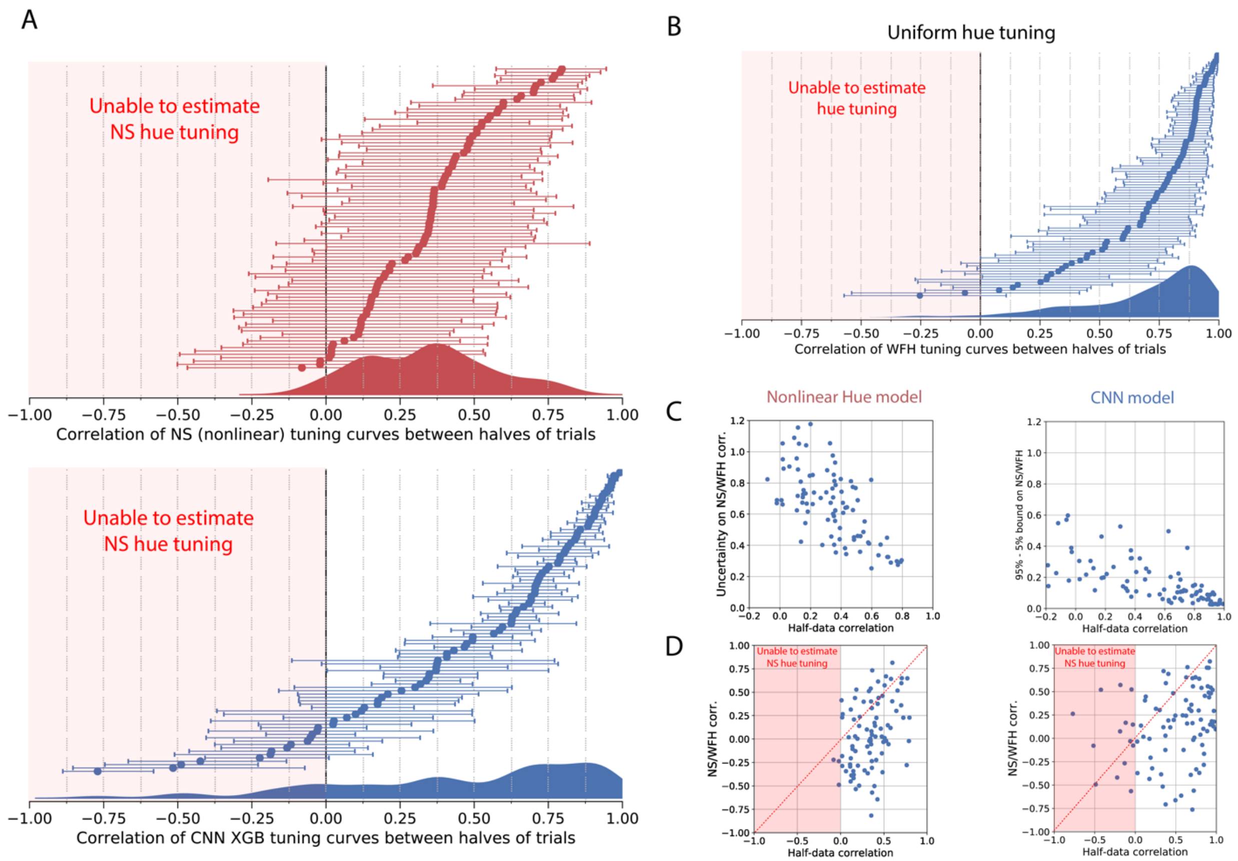


**Figure S1.** Our ability to estimate hue tuning can be captured by the correlation of the tuning estimated on two non-overlapping halves of the trials. This correlation would be 1 in the no-noise or infinite-data condition. For a single neuron, the half-trial correlation represents an estimate of what natural scene/uniform hue tuning curve correlations we would observe if hue tuning did not change between conditions. (Note that by fitting models on only half of the data, the estimate of hue tuning is noisier than in the full-data tuning estimation. Our actual ability to measure hue tuning is thus better than communicated by this control. For this reason these plots show a *lower bound* of the correlation we would expect if hue tuning were the same across conditions.) **A**) For the nonlinear hue model (red) and the CNN model (blue), these plots display the correlation of the tuning estimated on two non-overlapping halves of the trials. One can also consider this test as running 2-fold cross-validation and comparing the tuning curves estimated on both splits of data. Like in the main analysis, we split the data such that all trials (fixations) on the same image were placed in the same fold of data. In this plot the neurons are again ordered by their correlation to produce a cumulative distribution (the order of neurons is not the same as in Figures 2C and 4A). Errors show 5$^{th}$ and 95$^{th}$ percentiles of this procedure repeated on the original data resampled with replacement. The smoothed distributions projected below are reproduced in Figures 2C and 4A. **B)** The half-trial control for the uniform hue condition. This communicates how precisely we can estimate uniform hue tuning. The errors again derive from repeating the cross-half correlation when resampling the trials and re-splitting the data in half. **C)** The estimation error as communicated by these half-data control captures the same sources of variability that were incorporated into the principle uncertainty measure of the correlation between tuning curves (e.g. Figure 2Bii). That uncertainty was measured by resampling the trials, then re-calculating and re-correlating the tuning curves. To demonstrate this, here we show the relation between the half-data correlation and the size of the uncertainty bars from the main figures (Figures 2C and 4A). As expected, there is a strong negative correlation. Higher half-data correlations for a neuron correspond to smaller bounds of the natural scene/uniform hue correlation. **D)** Here we compare, neuron-by-neuron, the relationship between the half-data correlation and the natural scene/uniform hue correlation. (Panel A only communicates the difference in overall distributions.**)** Importantly, there is little relation between the half-data correlation (i.e. our ability to estimate natural scene hue tuning) and the natural scene/uniform hue correlation (i.e. whether we observed that neuron to shift tuning). This shows when we observe a shift in hue tuning, it is not simply because for that neuron we poorly estimated the natural scene hue tuning. Another key takeaway is the number of neurons that lie in the region below the dotted red line, where the split-trial correlation is higher than the natural scene/uniform hue correlation. For both estimation methods (nonlinear hue and CNN model), significantly more neurons lie below this line than above it.

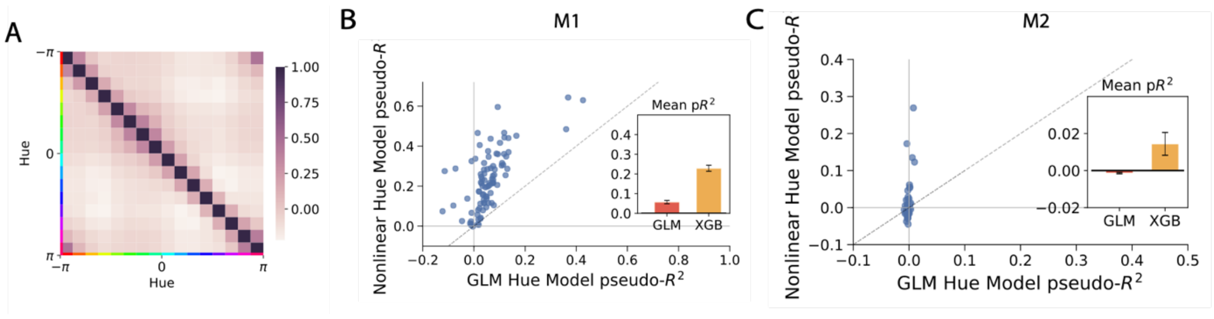


**Figure S2.** The response of V4 neurons to hue is nonlinear and contains interactions between bins of hues. A) The correlation matrix of hues on the natural image dataset observed by M1. Since the off-diagonal terms are not zero, there are correlation between hues (especially of similar colors). These correlations could bias the tuning curve of a linear fit if nonlinear hue interactions exist in the neural response. As shown in (B) and in (C), these interactions do indeed exist. This can be seen by the fact that the nonlinear model (gradient boosted trees, XGB) predicts neural activity better than the generalized linear model (GLM) when both are fed the (saturation-weighted) histograms of hues present within the receptive field during each fixation.

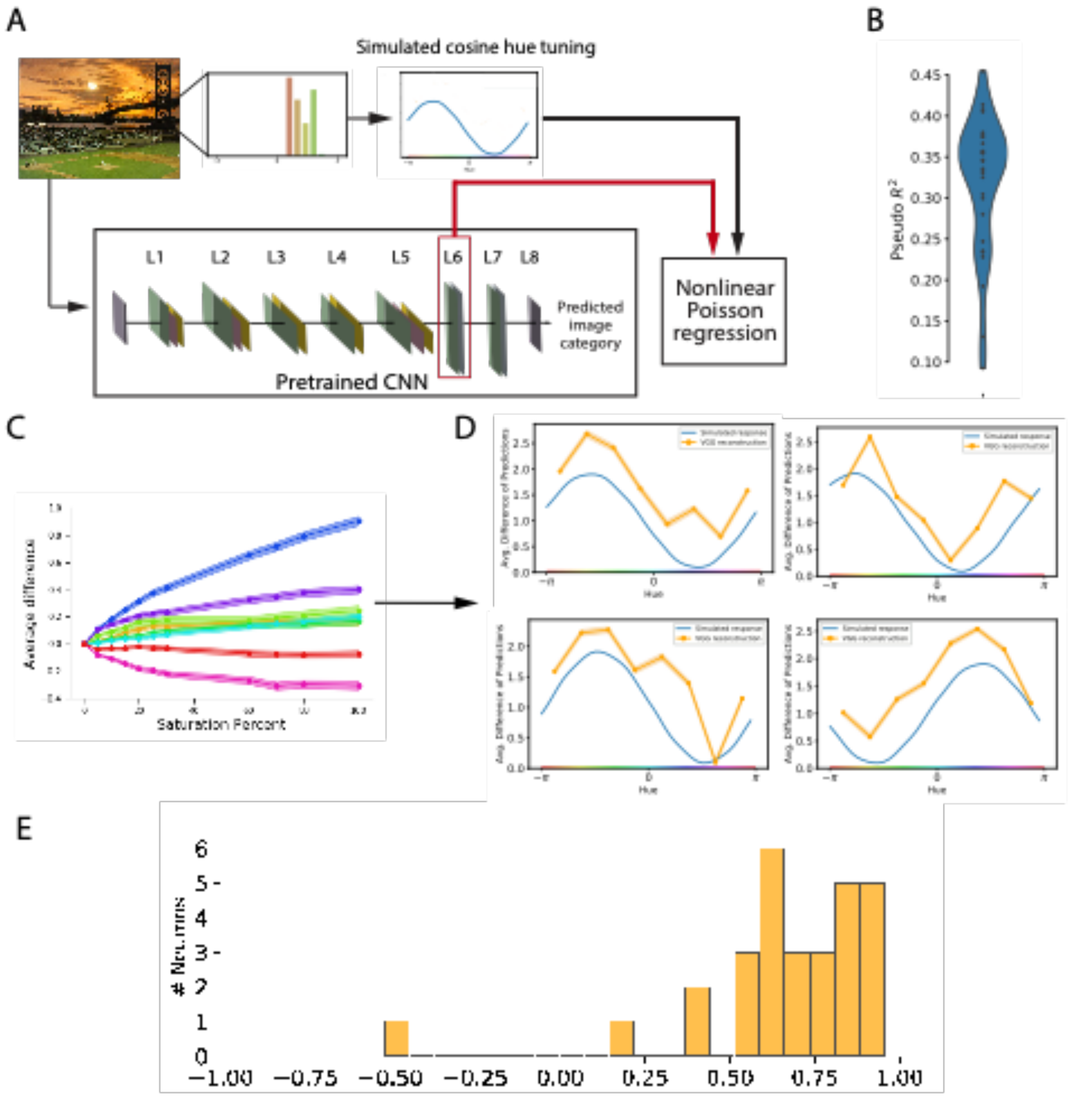


**Figure S3.** Reconstructing simulated neural responses shows that the CNN method can in principle observe hue tuning. A) As for the main model, we fit a nonlinear Poisson regression model to predict 'neural' responses from the intermediate activations of the VGG16 model when given image segments. Instead of the actual neural response, here we fit to a simulated neural response, which comprised of a random (fixed) cosine filter applied to the distribution of hues in each fixation. B) The model could predict these simple responses well, but not perfectly, with typical pseudo-$R^2$ scores less than 0.4. C) Once again we calculated the average difference in predictions between held-out images and those same images but with each of 8 bins of hues desaturated by some percentage. We plot the difference in response as a function of desaturation. It can be seen that the line is somewhat sub-linear, like for actual neural responses. This plot proves that some of this sub-linearity is not neural in origin, but rather a function of both our choice of color space (CIELUV) and the way that the VGG model incorporates color into the response. D) Tuning curves constructed in this way (from the slopes of the saturation dependencies) closely resemble the original filters, with some noise. E) The typical noise of this method's reconstruction of tuning curves can be summarized as a distribution of tuning curve correlations. This distribution is the point of comparison, representing what distribution we would expect if hue tuning were unchanged between categories of stimuli.

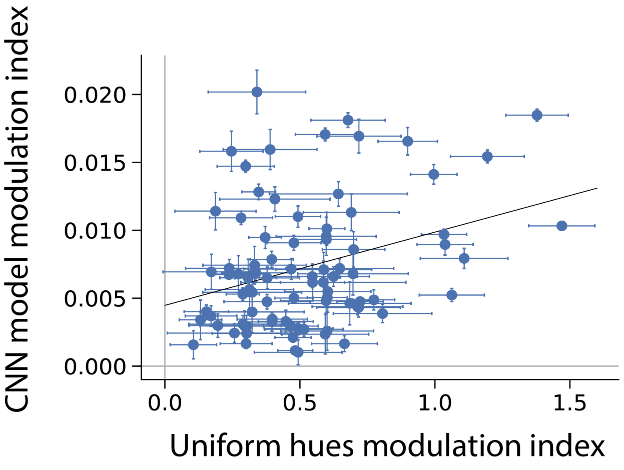


**Figure S4:** We calculated a modulation index measuring how drastically hue affected the V4 response on either uniform hues or natural images. In uniform hues, the modulation index was defined as the maximum of the uniform hue tuning curve, minus the minimum, and divided by the mean spike rate. In natural scenes, we examined how strongly various hues affected the CNN model response. This was measured by the difference between the maximum and the minimum of the CNN model tuning curve, which, measuring a difference in the predictions rather than the absolute value, is already mean-normalized. There was a weak correlation (p=0.003) between these two indices.
